# Lack of evidence supporting transgenerational effects of non-transmitted paternal alleles on the murine transcriptome

**DOI:** 10.1101/2022.12.23.521797

**Authors:** Rodrigo Gularte-Mérida, Carole Charlier, Michel Georges

## Abstract

Transgenerational genetic effects are defined as the effects of untransmitted parental alleles on the phenotype of their offspring. Well-known transgenerational genetic effects, in humans and other mammals, are the effects of a parental genotype on the nurturing ability of the parents, coined “genetic nurture”. However, there exist examples of transgenerational genetic effects in model organisms that are independent of nurturing effects and support the epigenetic transmission of a memory of the parental genotype possibly mediated by small RNA species. To test whether such transgenerational epigenetic effects might exist in mammals, we generated 833 isogenic C57BL/6J (B6) mice that differed only by the presence in the genome of their sire of one copy of four A/J chromosomes (MMU 15, 17, 19 or X). We measured 25 anatomical traits and performed RNA-Seq on five distinct tissues (heart, liver, pituitary, whole embryo, and placenta). There was no evidence of a significant effect from untransmitted A/J sire chromosome alleles, whether on anatomical traits or gene expression level. We observed an effect on *Mid1* expression levels in multiple tissues, but this was shown to be due to a de novo mutation that occurred in one of the sire lines. We conclude that transgenerational epigenetic memory of non-transmitted paternal alleles - if it exists - is uncommon in mice and likely other mammals.

## Introduction

Traits are typically considered to be genetic if individual phenotypes correlate with individual genotypes. In several instances, individual phenotypes also correlate with parental genotypes including with the non-transmitted alleles. These transgenerational genetic effects (TGE) include instances of “genetic nurture”, i.e., situations where the parental genotype will affect the environment in which the offspring is raised. TGE aim to identify parental genotypes and differ from transgenerational epigenetics where molecular changes are the result of environmental perturbations in the absence of genetic variation. This is easily understood by so-called “maternal effects”, which are determined by the care that is provided pre- and postnatally to offspring by mothers. Similar nurture effects of non-transmitted paternal alleles have been reported in humans, where the genotype of the parents may also affect the offspring’s environment (Mott et al. 2014, Kong et al. 2018; Cheuquemán, et al. 2021). In addition, the effects of non-transmitted parental alleles on the offspring’s phenotype exist that cannot be explained by genetic nurture. One such example is paramutation, which involves the epigenetic conversion of transmitted by non-transmitted alleles. Paramutation-like phenomena have been reported in plants, *D. melanogaster*, and possibly the mouse (Nadeau 2009; Heard and Martienssen 2014). A second example is the effect of gene copy number of *Sly, Ssty1, Ssty2, Srsy, Rbmy*, and *Rbm31y* of MMU Y on the predisposition and severity of experimental autoimmune encephalomyelitis in the next generation (Teuscher et al. 2006; Case et al. 2015). In C. elegans, the longevity of *Wd5*^*+/+*^ wild-type descendants of *Wd5*^*-/-*^ knockout worms was increased for three generations even in the absence of transmission of the *Wd5*^*-*^ knockout allele, when compared to worms of a fully *Wd5*^*+/+*^wild-type lineage (Greer et al. 2011). Most recently, promoter associated CpG DNA methylation via genetic engineering has been shown to be transmitted across multiple generations and retain parental phenotypic differences (Takahashi, et al. 2023). What remains unknown, however, is how common such nurture independent TGE are in mammals under naturally occurring variation.

We took advantage of mouse consomic strains and global gene expression by RNA-Seq to address this issue. Consomic mice, also referred to as chromosome substitution strains, harbour one chromosome pair from a donor strain “A” in the background of a recipient strain “B”. A genome-wide panel of consomic mice is commercially available for which “A” alleles from *A/J* mice were introgressed into the “B” background of *B6* mice (Nadeau et al. 2000). By backcrossing consomic males (say with a donor chromosome 15 of “A” ancestry) for two subsequent generations to the recipient “B” background strain, and selecting generation N2 offspring that have inherited a non-recombinant “B” chromosome 15, one obtains mice that are genetically identical to the inbred “B” background strain (isogenic), except for the genotype of their sire (heterozygous A/B for chromosome 15) (**Figure 1**). This design permits aggregation of any potential TGE across a whole chromosome, maximizing our chances to detect such effects if they exist. To study the effect of the non-transmitted paternal chromosome on many phenotypes at once, we decided to study growth and body composition traits in adult mice, and the transcriptome of five tissues at two developmental stages of the resulting N2 offspring by RNA-Seq.

**Figure 1.**
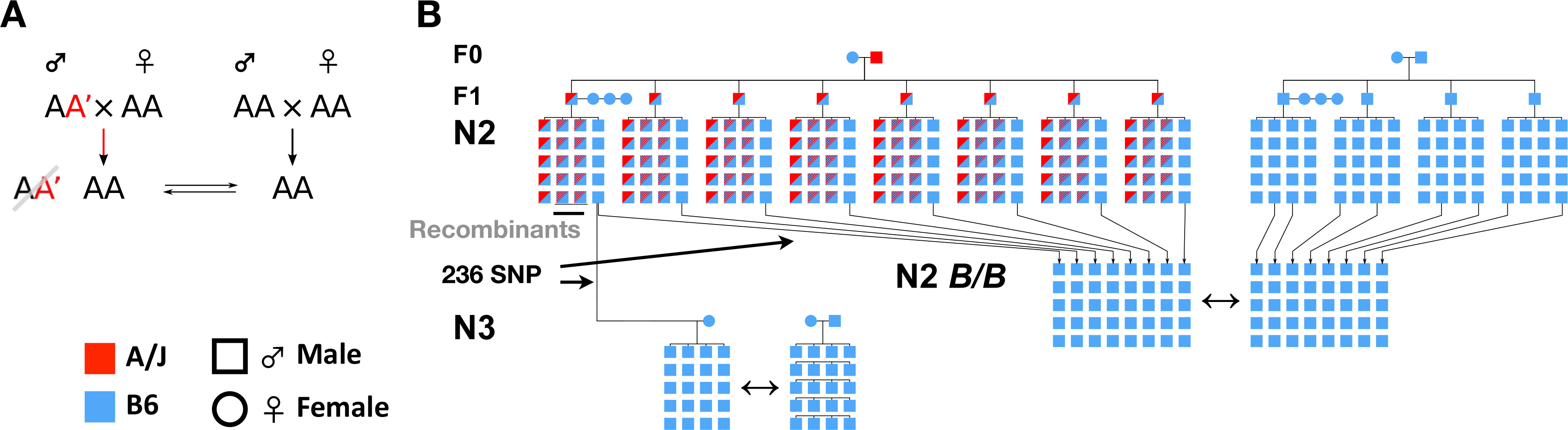
**A)** Single-locus two-allele model illustration of the mating design used to perform a test of hypothesis for paternal transgenerational genetic effects in isogenic mice. Where males heterozygous AA’ are mated to AA females. Resulting offspring are genotyped and only those with an AA genotype are compared to a control offspring on the right. **B)** Schematic representation of the cross performed for each B6.A/J–<N> CSS. Cross begins at F0 where a single male homozygous for each of chromosomes 15, 17, 19, and X, was mated to a single B6 female, and a single B6 control male is mated in parallel. The resulting males from the F1 were subsequently backcrossed with multiple B6 females, and N2 offspring genotyped for 236 SNP polymorphic between A/J and B6 to identify isogenic mice that inherit a complete B6 chromosome. Isogenic B6.A-<N> N2 ^*B6/B6*^ are then compared to the control contemporary group of B6/B6 purebred mice. A persistence N3 cross was performed by selecting an isogenic B6.A-<N> N2 ^*B6/B6*^ male mice and mated to multiple B6/B6 females and compared to a contemporary group of B6/B6 males. The only exception was B6.A-X in which B6.A-X F1 ^*B6/B6*^ isogenic mice are obtained within a single generation.

## Results

We performed this experiment for four of the 21 murine chromosomes taking advantage of the chromosome B6.A/J chromosome substitution strains (CSS) (MMU 15, 17, 19 and X. Hereinafter mouse strains will be denoted by their standard congenic nomenclature B6.A-<chr>, where B6 is the recipient strain, A is the A/J donor strain, chr refers to the specific mouse chromosome. Autosomes MMU 15, 17, and 19 were selected on the basis that MMU 15 was found the most transcriptionally diverse chromosome relative to B6, MMU 17 is known to affect body weight and body composition in other strains (Shockley and Churchill 2006, Corva et al. 2001; Farber et al. 2006), MMU 19 because it has a short chromosome length, and finally, MMU X was chosen for its convenience as one generation would automatically generate isogenic B6/B6 males. We produced 2,456 N2 mice, resulting in 958 consomic-derived *C57BL/6J* isogenic males throughout the experiment. The initial isogenic cohort was composed of 123 B6.A-15 N2 ^*B6/B6*^, 132 B6.A-17 N2 ^*B6/B6*^, 136 B6.A-19 N2 ^*B6/B6*^, 220 B6.A-X F1 ^*B6/B6*^, and 206 purebred *C57BL/6J* male controls. The corresponding proportions of recombinant and non-recombinant mice matched expectations based on chromosome size (in centimorgan), and were concordant with the observed results of Nadeau et al. (2000) (**Supplemental Material, Supplemental Table S1**). 80% of N2 male mice were euthanized at 60 days post-natal (dpn), dissected (recording 25 phenotypes pertaining to body weights, and body composition), and major organs flash-frozen – namely heart, liver, and pituitary. The other 20% of animals were collected in utero at 13.5 days post-coitum (dpc), embryo and placenta severed, and flash frozen. For each of the 10-line x timepoint combinations – (B6.A-15, B6.A-17, B6.A-19, B6.A-X, and B6 controls) x (60 dpn and 13 dpc) –, we extracted RNA from eight randomly selected mice within 1 standard deviation (SD) of the phenotypic mean for their corresponding genotypic group (60 dpn: heart, liver, and pituitary; 13 dpc: embryo, placenta). We then generated two equimolar RNA pools of four individuals for the 25-line x time point x tissue combinations, generating 50 libraries for next-generation RNA sequencing using TrueSeq® Stranded Total RNA with RiboZero kits (Illumina), and produced an average of 34.8 ± 8 million 100bp paired-end reads per library on an Illumina HiSeq 2000. One may expect the B6 background genome of the five lines to have diverged by the accumulation of *de-novo* mutations (DNMs) since their 19-year divergence. Such DNMs may cause differences in gene expression that may erroneously be attributed to TGE. To recognize such direct genetic effects, we resequenced the genomes of the founder male of each strain to 10-fold depth on an Illumina HiSeq2000 instrument. We identified a total of 2,794 non-A/J single nucleotide polymorphisms, 1,798 small insertion-deletions, and 63 copy number variants within the B6 genome of CSS founder males. The B6 genome of any two founders differed on average for 1,464 variants (**Supplemental Material**; **Supplementary Figure S1, Supplementary Table S2**).

RNA-Seq reads were mapped to the murine GRCm38 reference genome using the STAR aligner (Dobin et al. 2012; Engström et al. 2013). Read quantification and differential expression analysis between consomic-derived and *C57BL/6J* control samples (matched for tissue type) were conducted using HTSeq and DESeq2, respectively (Love et al. 2014; Anders et al. 2015). In a first analysis, we obtained 53 instances of significant differential expression (FDR 0.05) between consomic-derived and control samples involving 47 genes (**Figure 2A; Table 1**). Thirty-five (66 %) corresponded to increased expression in the consomic-derived samples (average fold change: 1.69; range: 0.228 – 4.33), and 18 (34%) to decreased expression (average fold change: -1.42; range: -0.274 – -3.68) (**Supplementary Figure S2**). The 53 instances were distributed across lines and sample types as shown in **Figure 2B**. Gene expressions were requantified using reverse transcriptase quantitative PCR assays (RT-qPCR) for the corresponding 47 genes in triplicate for all 200 samples that constituted the 50 RNA pools using the BioMarkHD qPCR system, Fluidigm at IntegraGen, Évry, France. Differential expression was confirmed (FDR ≤ 0.05) in at least one tissue for 10 of the 47 analysed genes. *Mid1, Crem* and *Gm26448* had the strongest evidence of a transgenerational genetic effect (technical replicates in **Figure 2A**; **Supplementary Table S3**).

**Table 1.** Number of significant differentially expressed genes in discovery and validation data sets.

| Discovery Population (N2.I) | Technical Replicate (N2.I) | Biological Replicate (N2.I) | Independent / Replicate Backcross (N2.III) | Persistence Backcross (N3) |
| --- | --- | --- | --- | --- |
| RNA-Seq<br>2 pooled libs<br>( N = 4 / lib; 50 libs ) | qPCR<br>8 per group<br>( N = 200 ) | qPCR<br>~20 per group<br>( N = 332 ) | qPCR<br>~20 per group<br>( N = 121 ) | qPCR<br>~20 per group<br>( N = 185 ) |
| 53 DE genes<br>FDR $\leq 0.05$ | 12 genes<br>$p \leq 0.05$ | 2 genes<br>$p \leq 0.05$ | 2 genes<br>$p \leq 0.1$ | 0 genes<br>$p \leq 0.05$ |
| | 1 gene<br>$p < 0.0001$ | 1 gene<br>$p < 0.0001$ | 1 gene<br>$p < 0.0001$ | 1 gene<br>$p < 0.0001$ |

**Figure 2.**
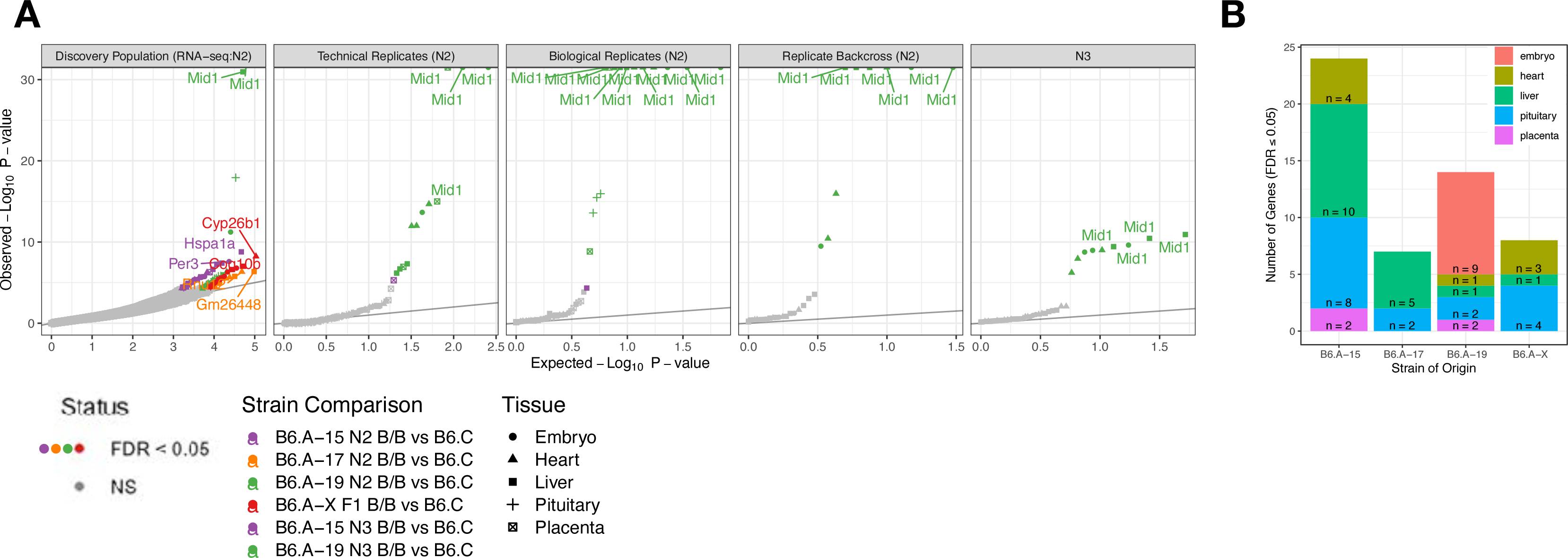
Differential expression analysis of RNA-Seq across five tissues comparing B6.A-<N> ^*B6/B6*^ and B6 controls. **A)** QQ-plots differential expression -log_10_ (*p*-values) of all genes compared across three cohorts, the Original N2 backcross (2012), a second independent Replicate N2 backcross (2013), and a persistence backcross N3 (2013). Within the N2 backcross, differential expression was measured three times. RNA-Seq of 50 libraries each composed of 4 mouse tissues, qRT-PCR of the same 200 mice for 53 genes with FDR ≤ 0.05 genes from RNA-Seq, and qRT-PCR of technically replicating genes on isogenic littermates from the same cohort. The latter set of technically replicating genes was measured in the Replicate N2, and Persistence N3 backcrosses. Colours represent the sire lineage. *P*-values were calculated by comparing each B6.A-<N> N2 ^*B6/B6*^ isogenic lineage to B6 Controls. Except for *Mid1*, from left to right our p-values show a loss of replication based on the significance and the directionality of the effects across all differentially expressed genes. **B)** Number of genes significant per B6.A–<N> N2 ^*B6/B6*^ vs B6.C comparison across the five tested tissues for each of the four tested CSS. B6.A-15 N2 ^*B6/B6*^ vs B6.C had the highest amount of differentially expressed genes, particularly in the liver. B6.A-19 N2 ^*B6/B6*^ vs B6.C had the second, totalling 14 genes, of which *Mid1* was consistently differentially expressed across all 5 tissues.

To further confirm the results obtained from the 10 remaining genes, we generated an independent replicate cohort with >500 male mice using the same mating design as above. The independent replicate cohort yielded 244 isogenic consomic-derived N2 males (MMU 17: 38; MMU 19: 19; MMU X: 98) and 89 *C57BL/6J* controls that were euthanized and dissected at 60 dpn. We extracted RNA from relevant tissue samples for an additional 14 to 26 animals of the discovery cohort, and 14 to 28 animals of the independent replicate cohort and evaluated the expression of the 10 candidate genes by RT-qPCR on the BioMarkHD using previously designed assays. Initial findings were confirmed in the independent samples from the discovery and replication cohort for one gene only: *Mid1* (**Figure 2A, Supplementary Table S3**). *Mid1* was highly significantly overexpressed (p < 0.0001; Median Log_2_ Fold change: 1.26 ± 0.4; range: 0.81 – 2.69) in all tissues of the MMU 19 consomic-derived animals (**Supplementary Figure S3**).

The *Mid1* gene (for *Midline 1*) encodes a member of the tripartite motif (TRIM) family that forms homodimers that associate with microtubules. Mutations in *Mid1* cause Opitz syndrome characterized by midline defects including cleft palate, laryngeal cleft, heart defects, hypospadias, and agenesis of the corpus callosum. In *Mus musculus domesticus, Mid1* maps to the X chromosome, yet its 5’ end is truncated by the pseudoautosomal boundary (PAB) on the Y chromosome (**Figure 3**). *Mid1* thus spans the PAB of *Mus musculus domesticus*. We examined the PAB in the five studied founder males and observed that the B6.A-19 founder had a duplication at the PAR that spanned Exon 4-10 of *Mid1* (**Figure 3**). The end of the duplication could not be precisely determined due to the limitations of the reference assembly (Morgan et al. 2019). Furthermore, our results support the hypothesis that the duplication is located on MMU Y, as all F1, N2, and N3 male offspring from this male inherited the X chromosome of B6 females for three generations, and the Y chromosome from the male. Thus, the differential expression of *Mid1* in the B6.A-19 consomic-derived isogenic mice when compared to the four other lines including controls is most likely due to the transmission of a duplicated *Mid1* transcript throughout the male lineage of the cross rather than to a TGE (**Supplementary Figure S4**). *Crem*, and *Gm26448* failed to replicate their initial effect through the experimental validations.

**Figure 3.**
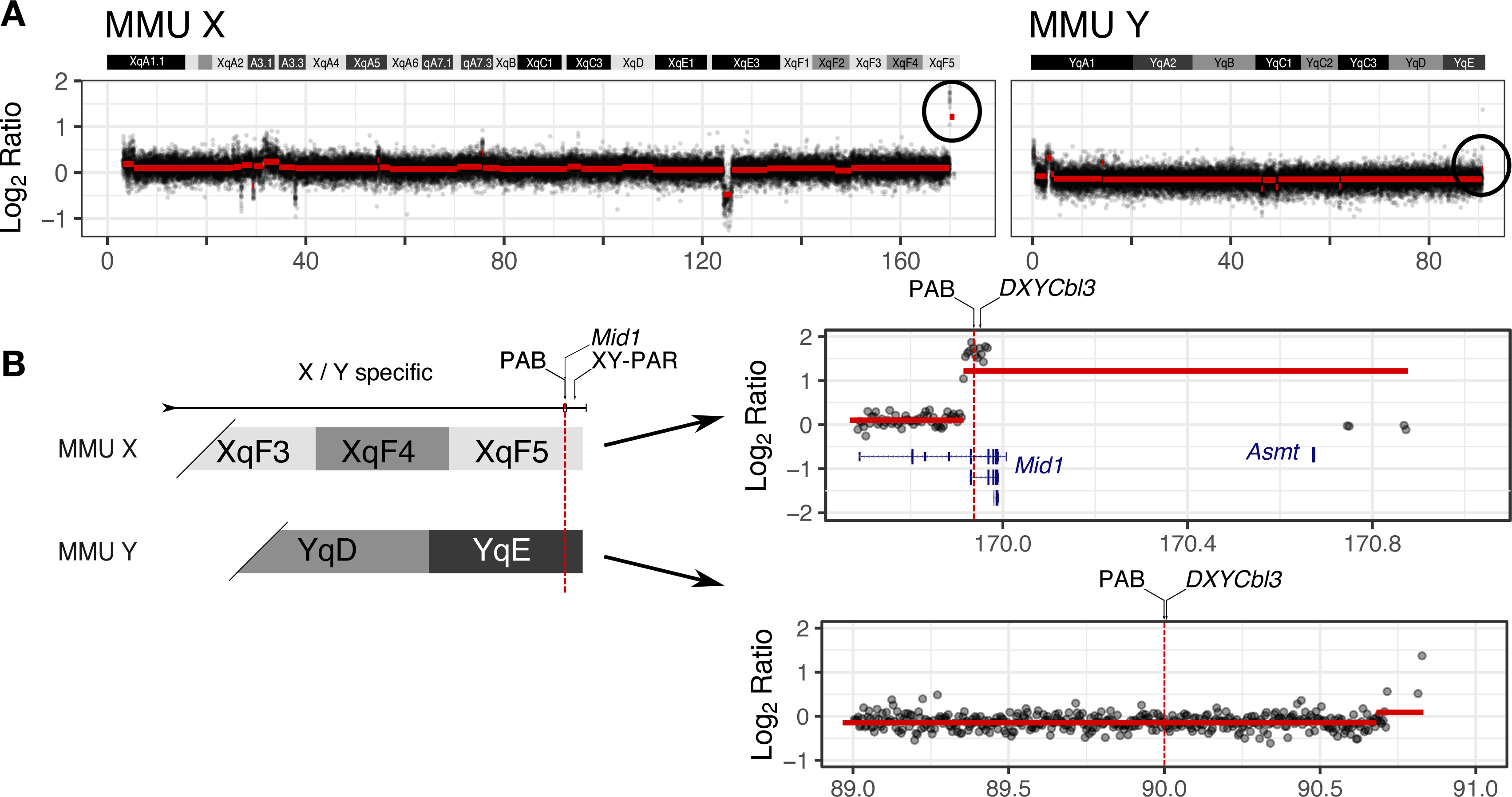
**A)** Segmented copy number profiles from the sex chromosomes of the B6.A-19 F0 founder male. At the distal end of mouse chromosome (MMU) X – left panel–, results show a duplication on the XY pseudo-autosomal region (PAR) highlighted with a circle. However, no duplication is observed in the MMU Y PAR highlighted in a circle (right panel). **B)** A closer view of the distal ends of MMU X and Y pseudo-autosomal boundaries (PAB), reveals the duplication event spans the *Mid1* gene, explaining the significant differential over-expression of B6.A-19 N2 ^*B6/B6*^ isogenic mice when compared to their B6 contemporary group.

We measured 25 anatomical phenotypes on the 1,370 60 dpn N2 animals from both the discovery and replication cohorts. We tested for possible TGE due to the non-transmitted paternal A/J chromosomes by comparing trait means between consomic-derived and control lines in two successive generations N2 and N3. There was no evidence of significant differences when accounting for multiple testing (*p*_*Bonf*_ *≤ 0*.*05;* **Supplementary Table S4**; **Supplementary Figure S5**). There was – at best – suggestive evidence (*p* = 0.0025) for a 6.7% reduction in heart weight in the B6.A-17 N2 consomic-derived line. This effect did not persist to the next generation through successive backcrossing, B6.A-17 N3 (**Supplementary Figure S6**).

## Discussion

Our work tested the hypotheses for genetically driven transgenerational effect via the paternal lineage comparing 25 anatomical traits of 1,164 isogenic male mice, and transcriptome-wide expression from five tissues – heart, liver, pituitary, embryo, and placenta. Our initial results suggested 3 genes *—Mid1, Crem*, and *Gm26448—* as differentially expressed between an isogenic-derived cohort relative to B6 purebred mice. Further validation of these genes, showed *Mid1*expression was differentially expressed across all B6.A-19 N2 ^*B6/B6*^, and B6.A-19 N3^*B6/B6*^ cohorts as a result of a segmental duplication at the XY-pseudoautosomal region (**Supplementary Figure S4**). All other 9 tested genes (including *Crem*, and *Gm26448*) failed to achieve significance in subsequent rounds of validation. We identified suggestive evidence for a 6.7% reduction in heart weight in the B6.A-17 N2^*B6/B6*^ consomic-derived line relative to B6.C purebred (**Supplementary Figure S5, Supplementary Figure S6**). Based on these results, we conclude that nurture independent TGE, if they exist at all, are extremely rare in mammals under naturally occurring genetic variation and are unlikely to contribute significantly to the phenotypic variance.

## Methods

### Animal husbandry

The initial breeding population purchased from the Jackson Laboratories (Bar Harbor, Maine) consisted of one male breeder and one female breeder approximately 5 weeks of age for each of the four chromosome substitution strains listed above, and one male breeder with three female breeders for C57BL/6J. These mice were placed under quarantine and then brought into the Specific Pathogen Free (SPF) facility of GIGA Research. After a seven-day acclimation period, males, and females for each of the four strains were coupled to maintain and expand all the strains in-house. Breeding cages were set up with one male and up to three females in clear polysulfone filter top cages (Model: 1290D EURO-STANDARD TYPE III, Tecniplast S.p.A, Varese-Italy). Breeding cages were checked daily to identify new-born pups and their corresponding dam. Whenever possible gestating females were transferred to an independent polysulfone cage for rearing pups, as SPF practices were to maintain pre-weaning pups and rearing females together with their sire. The presence or absence of the sire during pre-weaning did not affect any tested phenotype. At 30 ± 1 day of age males and females were weaned, identified with a numbered ear tag, divided into groups of up to 5, and placed in clear polysulfone filter top cages (Model: 1284L EURO-STANDARD TYPE II L, Tecniplast S.p.A). During the weaning process, mice were also weighed and a 1 - 2 mm tail biopsy was collected for genotyping. Breeding cages were fed a Standard SDS RM3 E diet (Cat. No. 801066, Special Diets Services-Essex, UK), whereas non-breeding and non-lactating cages were fed a maintenance diet, Standard SDS RM1 E diet (Cat. No. 801002, Special Diets Services). Enrichment was provided to all cages in the form of a half-white cardboard frites box which also provides nesting material. Experimental male mice from all cohorts N2, N3 and B6.C were sequentially weaned and placed in cages standardised with 5 males irrespective of strain. All mice were housed in accordance with the European Union Directive 2010/63/EU and the FELASA Recommendations and Guidelines (Guillen 2012), and experimental protocols approved by the Institutional Animal Care and Use Committee of the Université de Liège.

#### Characterization of 25 growth and body composition traits in adult and embryo male mice

The complete population of 966 N2 male mice from Cohort I, 125 N3 male mice from Cohort III, and 501 N2 male mice from Cohort IV were weighed at 30, 40, 50, and 60 days (d) of age. At 60 ± 1.5 d, mice were harvested as described in Gularte-Mérida (2015). Mice were anaesthetised with Isoflurane, during which time nasal-anal and nasal-tail lengths were collected. Mice were euthanised by decapitation under anaesthesia. Trunk blood was collected in BD Microtainer tubes containing Dipotassium EDTA (Cat. No. 365973, BD-Belgium), subsequently spun at 8000 rpm for 4 minutes, plasma was transferred to a 2 mL screw cap tube and stored at -20C. The weight of femoral, gonadal, mesenteric, and retroperitoneal fat pads, spleen, kidneys, liver, heart, gastrocnemius muscle, whole brain, and carcass weights were recorded up to the nearest 0.001 g. Lastly, femur length was measured on the left femur from the ball of the femur to the base. A collection of 11 tissues were flash frozen from each mouse and stored at -80C, these were spleen, liver, kidneys, heart, muscle, gonadal fat, brain, hypothalamus (separated after weighing), and lastly adrenal glands and pituitary gland were frozen but not weighed. A tail biopsy was also collected at the end of the harvest and used for DNA extraction.

Mouse N2 foetuses and placentas from cohort II were collected at 14 dpc and weighted and flash frozen in liquid nitrogen. This was achieved by isolating each female when the coital plug was identified during a morning inspection, day 0.5. Thirteen days later, females were euthanised by cervical dislocation, their uterus extracted, rinsed with DPBS (Cat. No BE17-512F, Lonza-Verviers), and subsequently kept on a petri-dish containing DPBS to avoid dehydration while foetuses were harvested under a stereoscope. Each amniotic sac containing one foetus and its respective placenta was isolated from the uterus, rinsed under a stream of DPBS. The amniotic sac was transferred to a new petri dish, opened carefully to maintain the yolk sac intact, and gently tearing it off from the placenta. Foetuses were excised by severing the umbilical cord under the stereoscope using fine tweezers and taken out from the yolk sac, rinsed in PBS, placed on a ruler, photographed, and then weighed to the nearest 0.001 g. Subsequently, the right leg was gently severed from the body with fine point tweezers to be used for DNA extraction. Placentas were inspected for any remaining maternal tissue, further cleaning any maternal tissue if needed, and weighed to the nearest 0.001 g. Both embryo and placenta were subsequently flash-frozen for later use.

### Statistical analyses

Organismal phenotypes were classified into three categories, body weight traits (comprising weights (g) at 14, 30, 40, 50, and 60 dpc. d), length traits (comprising Nasal-Anal, Nasal-Tail, and femoral lengths (mm)), and body composition traits (comprising liver, spleen, heart, kidney, brain, gastrocnemius muscle, carcass, and the four fat pad weights). To test for paternal transgenerational genetic effects the specific contrasts between each “test” group of B6.A-N N2^B6/B6^ isogenic derived mice and B6.C were analysed using general linear hypothesis testing with the R/*multcomp* package via the *glht* function set with a Tukey test (Bretz et al. 2010). Confounding factors such as litter size, presence of sire in the pre-weaning cage, and dam’s age were tested as covariates during the model selection across all phenotypes. However, none of these showed any significant effects to change the conclusions of the analyses and thus were excluded from the final comparisons. A summary of power calculations is presented in **Supplementary Material**.

### Characterization of global gene expression by RNA-Seq of adult and embryonic tissues

To characterize and analyse global gene expression we extracted RNA from 8 individuals in three adult tissues — liver, heart, pituitary —, whole embryo and placenta in each of the five isogenic derived sub-cohorts (B6.A-15 N2 ^B6/B6^, B6.A-17 N2 ^B6/B6^, B6.A-19 N2 ^B6/B6^, B6.A-X F1 ^B6/B6^, and B6.C) for a total of 200 samples. Libraries were constructed using Illumina TrueSeq® Stranded Total RNA with RiboZero kit (No. Cat. RS-122-2201 or -2202). We constructed two pooled libraries for each tissue × strain combination from equimolar quantities of RNA of four individuals. RNA-Seq libraries were sequenced on an Illumina HiSeq 2000 where each lane contained five libraries, one per strain.

We obtained on average 32.5 M reads per library, a total of 162.5 M reads per lane. Sequence reads were aligned to the mouse reference assembly GRCm38 with alternative loci using STAR aligner (Dobin et al. 2012) using the annotated 2-pass method (annotation with Ensembl Genes 75) described by Engström et al. (2013), where « splice junctions » are obtained from the first pass, and the second pass refines the alignment of all sequence reads. Following the alignment, we analysed sequences to 1) detect differential expression in the five cohorts of isogenic mice; and 2) identify if genetic variation exists among the strains that may explain the differences in significantly differentially expressed genes (details in **Supplementary Material**). For the first analysis, we used HTSeq to quantify the expression of each gene in the Ensembl 86 annotation and we used DESeq2 to perform the differential expression analyses of all genes (Love et al. 2014). Comparisons were made as described in the previous section where each isogenic derived sub-cohort (B6.A-15 N2 ^*B6/B6*^, B6.A-17 N2 ^*B6/B6*^, B6.A-19 N2 ^*B6/B6*^, B6.A-X F1 ^*B6/B6*^) is compared to B6.C. At a Log2FoldChange of ≥ 2, power of detection using the 2 pooled RNA-Seq samples ranged from ≥ 0.90 in the adult tissues(pituitary, liver, and heart), to 0.6 for the embryo and placenta (**Supplementary Material**).

### Technical and biological replications of RNA-Seq differentially expressed genes by RT-qPCR

To confirm our initial findings, we carried out a technical replication on the same 200 samples used to construct the RNA-Seq libraries. In these samples, we re-measured the expression level of the 48 genes found differentially expressed by RNA-Seq using reverse transcriptase quantitative PCR (RT-qPCR) in triplicate determinations using Fluidigm’s BioMarkHD qPCR system. This platform allows us to simultaneously test up to 32 assays in triplicate in 96 samples (a 96 × 96 factorial design on a single reaction plate), for a total of 9,216 RT-qPCR reactions per plate, and each gene was measured in two or more tissues. Primer assays for the RT-qPCR of the technical replicate and first validation cohorts were designed by IntegraGen (Évry, France) (**Supplementary Table S5**). For the top-ranking differentially expressed genes –*Mid1, Coq10b, Per3, Ide, and Crem*– we designed multiple assays targeting different exon-exon junctions maximising the number of transcript variants assayed. For the remaining 43 genes, one assay per gene was designed (**Supplementary Table S3**). In addition, we amplified eight genes used routinely as endogenous controls, *B2m, Eif3j1, Gusb, Hprt, Rpl37rt, Rp10, Tbp*, and *Tubb5* each with one primer assay. Complementary DNA (cDNA) was synthesized from RNA using Superscript III following the manufacturer’s recommendations (Life Technologies, Carlsbad, CA). A reference sample was created by pooling cDNA from all the samples of each tissue, from which a 1:4 serial dilution was carried out to develop a four-point standard curve. All reactions were carried in two stages, where the first stage cDNA is a pre-amplification using TaqMan PreAmp Master Mix (Cat No. 4384267) following the manufacturer’s recommendations, and the second stage is the RT-qPCR reaction on the BioMarkHD system using EvaGreen(R) master mix.

Differential expression was analysed using relative gene expression based on the 2^-Ct^ method (Livak and Schmittgen 2001). First, we perform endogenous control gene selection based on the geNorm algorithm (Vandesompele et al. 2002) using qBase+ (Biogazelle, Belgium). These analyses suggested the use of *Tubb5* and *Gusb* as endogenous controls in embryo, *Tbp* and *Hprt* in the placenta, liver, and pituitary, and all four genes, *Tubb5, Gusb, Tbp* and *Hprt* in the heart. Expression values estimated as 2^-Ct^ were calculated using the R/Bioconductor *ddCt* package (Zhang et al. 2015), where the median Ct for each sample is corrected by the median of the pooled reference sample, and to the median of the endogenous control genes as described in Livak & Schmittgen (2001).

The 14 genes replicated between RNA-Seq and the RT-qPCR differential expression –*Clpx, Srebf1, Col1a1, Crem, Gm129, Gm26448, Hpsa1a, Hpsa1b, Hspg2, Mid1, Per3, Tef*, and *Thrsp–. These* were quantitated in an additional 800 N2 littermates from all the cohorts developed throughout the course of the experiment. Roughly 20 mice for each of the four B6.A-15 N2 ^*B6/*B6^, B6.A-17 N2 ^*B6/B6*^, B6.A-19 N2 ^*B6/B6*^, and B6.C isogenic littermates (Cohort I and II), 20 mice from each of B6.A-17 N3 ^*B6/B6*^, B6.A-19 N3 ^*B6/B6*^, and B6.C N3 controls (Cohort III), and 20 mice from each of B6.A-17 N2 ^*B6/B6*^, B6.A-19 N2 ^*B6/B6*^, and B6.C independent replicate backcross (Cohort IV). The cDNA synthesis reactions, RT-qPCR technical replicate, and first validation results were performed by IntegraGen (Évry, France), using *Tbp, Hprt, Tubb5*, and *Gusb* as endogenous control genes. Expression values for this first validation were estimated as previously described and analysed by comparing the expression values from the different B6.A-<N> N2 ^*B6/B6*^ to B6.C as described in the statistical analysis section.

Based on the gene expression results from the previous RT-qPCR expression analyses, a second of validation was performed on *Crem* and *Gm26448* in the pituitary. These two genes and corresponding endogenous controls were measured in 32 B6.A-15 N2 ^*B6/B6*^, and 17 B6.A-17 N2 ^*B6/B6*^ and 23 B6.C new male littermates from Cohort I. Also, included were an additional 14 B6.A-17 N2 ^B6/B6^ and 27 B6.C new littermates from Cohort IV. Complementary DNA from these samples was synthesised using SuperScript IV following manufacturer’s recommendations. Where *Crem, Tbp* and *Hprt* had pre-designed assays from IDT PrimeTime(R) probes and these were chosen to measure their expression, whereas *Gm26448* was measured with a custom IDT PrimeTime(R) assay (**Supplementary Table S3**). Real-Time qPCR reactions were carried out according to manufacturer’s recommendations. Briefly, each reaction was composed of 1X PrimeTime(R) Universal Master Mix (Cat No. 1055772, 0.2 μM PrimeTime(R), and 10ng of cDNA.

## Data availability

Sequence data for the mouse genomes and isogenic transcriptomes will be available through the European Nucleotide Archive project (PRJEB70518). Genotype and phenotype data will be publicly accessible at http://www.github.com/RodrigoGM/tgv_2023/

## Acknowledgments

This project was funded by a grant from the Walloon Ministry of Agriculture (DGARNE). We are grateful to all the personnel of the GIGA Mouse Housing and Transgenics Platform, in particular Benoit Remy, Audrey Tromme, and Fabien Ectors. We also give special thanks to the members of the GIGA Genomics Platform Nadine Cambisano, Naïma Ahariz, and Wouter Coppieters for their assistance in sample processing.

## Supplemental Information

### Supplemental Material

Extended methods for generating the backcross population, genotyping, selection of isogenic mice, power calculations, and sequence analysis of founder males

**<u>Supplementary Table S1.</u>** Differential expression analysis of isogenic B6/B6 derived mice vs B6 Controls.

**<u>Supplementary Table S2.</u>** Means ± SE for body weights, growth, body length, and body composition traits measured in isogenic B6/B6 N2 mice derived from sires with different genotypes.

**<u>Supplementary Table S3.</u>** Gene expression assay design to validate differentially expressed genes from RNA-Seq between B6.A-<N> and B6.C.

**<u>Supplementary Table S4.</u>** Summary of genotype and allelic frequencies across all 11 backcrosses.

**<u>Supplementary Table S5.</u>** DNA sequence polymorphisms and de-novo mutations in B6.A-15, B6.A-17, B6.A-19 and B6.A-X.

**<u>Supplemental Figure S1.</u>** The mutational and copy number landscapes of de-novo and A/J alleles across the B6.A-<N> founder males of the tested CSS.

**<u>Supplemental Figure S2.</u>** Volcano plots of differential expression across all derived isogenic N2 mice from the four CSS.

**<u>Supplemental Figure S3.</u>** Global expression and *Mid1* effects comparisons across replicates.

**<u>Supplemental Figure S4.</u>** Inheritance model of the *Mid1* segmental duplication across the B6-A-19 male lineage throughout the discovery, validation, and persistence experimental crosses.

**<u>Supplemental Figure S5.</u>** QQ-plots of -log_10_ *p*-values of all 25 anatomical traits across all three analyzed cohorts.

**<u>Supplemental Figure S6.</u>** Heart, and body weights weight across the Discovery (N2), Replicate (N2), and Persistence (N3) backcrosses in B6.A-17 N2 ^*B6/B6*^, B6.A-X F1 ^*B6/B6*^, and B6 Control mice.

## References

Anders S, Pyl PT, Huber W. 2015. HTSeq--a Python framework to work with highthroughput sequencing data. Bioinformatics 31: 166–169.

Case LK, Wall EH, Osmanski EE, Dragon JA, Saligrama N, Zachary JF, Lemos B, Blankenhorn EP, Teuscher C. 2015. Copy number variation in Y chromosome multicopy genes is linked to a paternal parent-of-origin effect on CNS autoimmune disease in female offspring. Genome Biology 16: 28–14.

Cheuquemán C, Maldonado R. 2021. Non-coding RNAs and chromatin: key epigenetic factors from spermatogenesis to transgenerational inheritance. Review. Biol Res. 2021 Dec 20;54(1):41.

Corva PM, Horvat S, Medrano JF. 2001. Quantitative trait loci affecting growth in high growth (hg) mice. Mammalian Genome 12: 284–290.

Dobin A, Davis CA, Schlesinger F, Drenkow J, Zaleski C, Jha S, Batut P, Chaisson M, Gingeras TR. 2012. STAR: ultrafast universal RNA-Seq aligner. Bioinformatics 29: 15–21.

Engström PG, Steijger T, Sipos B, Grant GR, Kahles A, Rätsch G, Goldman N, Hubbard TJ, Harrow J, Guigo R et al. 2013. Systematic evaluation of spliced alignment programs for RNA-seq data. nature methods 10: 1185–1191.

Farber CR, Corva PM, Medrano JF. 2006. Genome-wide isolation of growth and obesity QTL using mouse speed congenic strains. BMC Genomics 7: 102.

Gapp K, Jawaid A, Sarkies P, Bohacek J, Pelczar P, Prados J, Farinelli L, Miska EA, Mansuy IM. 2014. Implication of sperm RNAs in transgenerational inheritance of the effects of early trauma in mice. Nature Neuroscience doi:10.1038/nn.3695.

Greer EL, Maures TJ, Ucar D, Hauswirth AG, Mancini E, Lim JP, Benayoun BA, Shi Y, Brunet A. 2011. Transgenerational epigenetic inheritance of longevity in Caenorhabditis elegans. Nature 479: 365–371.

Heard E, Martienssen RA. 2014. Transgenerational epigenetic inheritance: myths and mechanisms. Cell 157: 95–109.

Kong A, Thorleifsson G, Frigge ML, Vilhjalmsson BJ, Young AI, Thorgeirsson TE, Benonisdottir S, Oddsson A, Halldorsson BV, Masson G et al. 2018. The nature of nurture: Effects of parental genotypes. Science 359: 424–428.

Livak KJ, Schmittgen TD. 2001. Analysis of relative gene expression data using real-time quantitative PCR and the 2(-Delta C(T)) Method. Methods 25: 402–408.

Love MI, Huber W, Anders S. 2014. Moderated estimation of fold change and dispersion for RNA-Seq data with DESeq2. In biorxivorg, doi:10.1101/002832.

Morgan AP, Bell TA, Crowley JJ, Pardo-Manuel de Villena F. 2019. Instability of the Pseudoautosomal Boundary in House Mice. Genetics 212: 469–487.

Mott R, Yuan W, Kaisaki P, Gan X, Cleak J, Edwards A, Baud A, Flint J. 2014. The Architecture of Parent-of-Origin Effects in Mice. Cell 156: 332–342.

Nadeau JH. 2009. Transgenerational genetic effects on phenotypic variation and disease risk. Human Molecular Genetics 18: R202.

Nadeau JH, Singer JB, Matin A, Lander ES. 2000. Analysing complex genetic traits with chromosome substitution strains. Nature Genetics 24: 221–225.

Shockley KR, Churchill GA. 2006. Gene expression analysis of mouse chromosome substitution strains. Mammalian Genome 17: 598–614.

Takahashi Y, Morales Valencia M, Yu Y, Ouchi Y, Takahashi K, Shokhirev MN, Lande K, Williams AE, Fresia C, Kurita M, Hishida T, Shojima K, Hatanaka F, Nuñez-Delicado E, Rodriguez Esteban C, Izpisua Belmonte JC. 2023. Transgenerational inheritance of acquired epigenetic signatures at CpG islands in mice. Cell 186, 715–731.

Teuscher C, Noubade R, Spach K, McElvany B, Bunn JY, Fillmore PD, Zachary JF, Blankenhorn EP. 2006. Evidence that the Y chromosome influences autoimmune disease in male and female mice. Proceedings of the National Academy of Sciences of the United States of America 103: 8024–8029.

Vandesompele J, De Preter K, Pattyn F, Poppe B, Van Roy N, De Paepe A, Speleman F. 2002. Accurate normalization of real-time quantitative RT-PCR data by geometric averaging of multiple internal control genes. Genome Biology 3: RESEARCH0034.

