## Supplemental Material for "Lack of evidence supporting transgenerational effects of non-transmitted paternal alleles on the murine transcriptome"

^1^Unit of Animal Genomics, GIGA Institute and Faculty of Veterinary Medicine, University of Liège, 1 Avenue de l’Hôpital, 4000 Liège, Belgium. ^2^Current address: Department of Surgery, Memorial Sloan Kettering Cancer Center, New York, NY

### Supplemental Material

#### Generating isogenic B6 mice from B6.A F1 consomic sires

To uncover TGE (Figure 1) in growth, body composition, and gene expression phenotypes we generated mice that were genetically identical ("isogenic”) for C57BL/6J (B6) genotype whose sire differed by the genotype for one complete chromosome, namely MMU 15, MMU 17, MMU 19 and MMU X, relative to a B6 control. To this end, "test" animals were B6 obtained by mating B6 dams with F1 "consomic” sires carrying one A/J chromosome in an otherwise B6 background, referred herein by their congenic nomenclature as B6.A-Chr N2; followed by their genotype A/B6, A/_, B6/B6 (where A is the A/J genotype) to denote heterozygous non-recombinant, recombinant, and B6 non-recombinant mice. The "control" animals were B6 purebred animals, born from purebred B6 sires and B6 dams, these are referred to as B6.C. When we refer collectively to the consomic strains of this experiment we denote them as B6.A-Chr. By genotyping for 236 SNP markers differentiating the B6 alleles from the A/J alleles across MMU 15, 17, 19, X, and Y, we selected offspring that inherited a non-recombinant B6 chromosome from their heterozygous sire recovering a complete B6 genome. The visual representation of the 236 SNP markers across all chromosomes is shown in Figure S1, and the resulting mouse genotypes of all backcrosses described below are shown in Figure S2.

Four sires homozygous for A/J alleles on chromosomes MMU 15, MMU 17, MMU 19 and MMU X in an otherwise B6 background, from the Jackson Laboratories (Bar Harbor, USA). Sires corresponded to the chromosome substitution strains C57BL/6J-Chr 15^A/J^/NaJ (Stock No: 004393), C57BL/6J-Chr 17^A/J^/NaJ (Stock No: 004395), C57BL/6J-Chr 19^A/J^/NaJ (Stock No: 004397) and C57BL/6J-Chr X^A/J^/NaJ (Stock No: 004398). These sires constituted our Fø generation and were mated to purebred C57BL/6J (Stock No: 000664) females (purchased from Charles River Laboratories, France) to yield four F1 populations that were uniformly heterozygous A/B6 for their corresponding chromosome. Subsequently, F1 sires were backcrossed to B6 females, to generate three segregating N2 populations, and one non-segregating N3 population. Only males were used to test TGE.

The first N2 backcross population (Cohort I) was produced between August 2011 and June 2012. It comprised 308 B6.A-15 N2, 209 B6.A-17 N2, 247 B6.A-19 N2 and 48 B6.A-X F1 male offspring, as well as 67 control male B6.C offspring. The number of F1 sires to generate the N2 was 15 B6.A-15 F1, 16 B6.A-17 F1, 22 B6.A-19 F1, one B6.A-X F1 and 10 B6.C. The number of litters per strain generated in this N2 population was 97 B6.A-15 N2, 55 B6.A-17 N2, 79 B6.A-19 N2, 14 B6.A-X F1 and 26 B6.C, with an average size (± SD) 6.9 ± 3.4 for B6.A-15 N2, 7.5 ± 3.4 for B6.A-17, 6.6 ± 3.0 for B6.A-19 N2, 6.0 ± 3.1 for B6.A-X F1 and 6.3 ± 3.0 for B6.C. The population of N2 males from cohort I were reared until the age of 60 days and then euthanised for tissue collection. Note that the MMU X derived subpopulations differ in generation. This is due that in the F1, male offspring inherit a copy of MMU Y from their sire, and chromosome X is inherited from their dam, thus only one backcross is required to generate B6/B6 non-recombinant males. However, we were concerned with the possibility of segregation at the XY Pseudo-Autosomal Region (XY-PAR), thus markers were genotyped in this region to identify males with recombination events in the XY-PAR.

The second backcross population (Cohort II) was produced between November and December 2012. It comprised 229 B6.A-15 N2, 229 B6.A-17 N2, 235 B6.A-19 N2 and 105 B6.A-X F1 offspring, as well as 59 B6.C. We used the same F1 sires from Cohort I, but purchased 200 B6 females 5 weeks of age (Charles River Laboratory, France). The average litter size in this cohort was of 8.1 ± 1.7 in B6.A-15 N2, 7.9 ± 1.9 in B6.A-17 N2, 7.9 ± 1.6 in B6.A-19 N2, 8.1 ± 1.4 in B6.A-X F1, and 8.1 ± 1.4 in B6.C controls. In this cohort, females gestations were interrupted at 14 days post-coitum (dpc) and the fetuses and placentas (males and females) were collected. Foetuses were genotyped for chromosomes X and Y to determine their gender and only male foetuses were used for further analysis.

The third population (Cohort III) was an N3 backcross produced between December 2012 and May 2013 to study the persistence of TGE by backcrossing isogenic B6 N2 males to B6 females. It comprised 25 B6.A-15 N3, 22 B6.A-17 N3, 29 B6.A-19 N3, and 23 B6.A-X N2 male offspring, as well as 20 B6.C. The number of N2 sires used to develop this cohort were five B6.A-15 N2, three B6.A-17 N2, four B6.A-19 N2, three B6.A-X F1, and two B6.C. The number of litters produced in the N3 population was 10 B6.A-15 N3, six B6.A-17 N3, seven B6.A-19 N3, seven B6.A-X N2, and three B6.C. The average litter size in this cohort was 5 ± 2 for B6.A-15 N3, 7 ± 3 for B6.A-17 N3, 8 ± 4 B6.A-19 N3, 7 ± 4 B6.A-X N2, and 12 ± 3 for B6 controls. N3 males from Cohort III were reared until the age of 60 days and then euthanised as those in Cohort I.

Finally, the fourth N2 backcross population (Cohort IV) was an independent replicate N2, between January and September 2013 to confirm the results obtained in Cohort I. We used independent homozygote males several generations apart from the original Fø reared in our breeding colony as the grandsires. Cohort IV comprised 232 B6.A-17 N2, 80 B6.A-19 N2, 107 B6.A-X F1 male offspring, as well as 82 purebred B6 control males. The number of F1 sires used was 24 B6.A-17 F1, 12 B6.A-19 F1, eight B6.A-X Fø, and nine purebred B6. The number of litters produced was 65 B6.A-17 N2, 26 B6.A-19 N2, 28 B6.A-X F1, and 19 purebred B6. The average litter size per strain was of 7.5 ± 2.7 for B6.A-17 N2, 6.2 ± 3.2 B6.A-19 N2, 6.7 ± 2.4 B6.A-X F1, and 7.3 ± 2.2 purebred B6.C. Males from Cohort IV were reared until the age of 60 days and then euthanised as done for those in Cohort I and III.

Analysis of the corresponding genotypes allowed us to sort the N2 populations in offspring that inherited (i) a non-recombinant A chromosome, (ii) a recombinant B6-A chromosome, and (iii) a non-recombinant B6 chromosome (Figure 1). The corresponding proportions did not deviate significantly between cohorts and averaged (i) 0.24, 0.22, 0.25; (ii) 0.49, 0.55, 0.49; and (iii) 0.26, 0.22, 0.24, for B6.A-15 N2, B6.A-17 N2, B6.A-19 N2, respectively. These proportions correspond to map lengths of 77 cM on MMU-15, 65 cM on MMU-17, and 59 on MMU-19.

#### SNP genotyping, recombination analysis, and identification of isogenic offspring in the three backcrosses

We designed two multiplex marker panels referred to as P1 and P2, comprising 234 and 242 SNP markers, respectively. The markers differentiated between B6 and A/J alleles and were distributed evenly across MMU 15, MMU 17, MMU 19, and the pseudo-autosomal region (PAR) (Figure S1). P2 is an extension of P1, containing additional markers on the X and Y chromosomes for sex determination of the foetuses. The panels were designed for SNP genotyping by mass spectrometry using Sequenom MassARRAY® (San Diego, CA), now Agena Biosciences, in collaboration with Neogen/GeneSeek (Lincoln, NE). Genomic DNA was extracted for each animal from a tail biopsy using Promega Maxwell Tissue cartridges (Cat. No. AS1610, Promega, Fitchburg, WI). We genotyped the ensemble of 1,467 adult male mice, 881 embryos of 14 dpc, and 200 additional individuals with known genotypes as controls (homozygote B6.C, known homozygote, and heterozygote mice for each of the four A/J consomic strain). Cohort I was genotyped with P1, while cohorts II and IV were genotyped with P2. Genotyping was carried out at Neogen/GeneSeek (Lincoln, NE). The analysis of these genotypes identified 72 B6.A-15 N2*^B6/B6^*, 49 B6.A-17 N2*^B6/B6^*, 55 B6.A-19 N2*^B6/B6^*, and 48 B6.A-X F1*^B6/B6^* apparently isogenic male mice. In the embryo backcross, we identified 46, 48, 59, 93 and 59 male foetuses of 14 dpc isogenic for B6.A-15 N2*^B6/B6^*, B6.A-17 N2*^B6/B6^*, B6.A-19 N2*^B6/B6^*, B6.A-X F1*^B6/B6^*; and B6.C, respectively. Lastly, in the second adult backcross, we identified 38, 19, 98, and 89 apparently isogenic mice in B6.A-17 N2*^B6/B6^*, B6.A-19 N2*^B6/B6^*, B6.A-X F1*^B6/B6^*, and B6.C respectively. All genotypes and phenotypes are provided as Supplemental Data.

We compared our observed number of recombinant and non-recombinant mice, and recombination rates with those of Nadeau et al. (2000), and the expected recombination rates of the new MGI standard genetic map from Cox et al. (2009) using a Fischer Exact test. Our results were highly concordant with those of Nadeau et al. (2000) (p > 0.1) for all chromosome crosses ([**Supplementary Table S1**](https://docs.google.com/spreadsheets/d/16Ig1gN1bj61P2uPjkO7YJd45vWzoLpbT/edit#gid=1107694650)). However, when compared to the new MGI standard genetic map, the estimates for recombination rates from our three independent MMU 19 back-crosses, and that of Nadeau et al. (2000) deviate significantly from the expected recombination rate (p ≤ 10^-6^); whereas all other chromosomes did not deviate from the expectation (p > 0.09). We suspect either an error in the estimates or additional mouse strains contributed to the results of Cox et al. (2009).

#### Experimental power and sample size

### Power and sample size estimates were performed for all anatomical traits using the n.ttest function in the R/samplesize package. For each trait, sample size per group was estimated to detect an four effect sizes of 5%, 10%, 15% and 20% of the trait mean, at 9 power levels (50%, 60%, 70%, 80%, 85%, 90%, 95%, 99%, and 99.9%) using trait specific mean and variances. With the exception of subtle changes (5%), and most body fat traits, an 80% power of detection was achieved in our comparisons (Figure S3). RNA-seq power of detection was performed using the R/ssizeRNA package across all five tissues. Power estimations suggest our approach is at the limit of detection (Table S1).

#### Discovery of DNA variation in the « background » of the chromosome substitution strains

### Our experimental design assumes that we are comparing a completely homozygous B6/B6 in the background genome after two generations of backcrossing to B6 purebred females (Figure 2). To investigate the possibility that a « strain-specific » mutation that arose during the chromosome substitution strain derivation or by the isolated breeding of the strains causes the differential expression we re-sequenced all Fø founding sires purchased from JAX®. One whole genome PCR-free library was constructed per male using Illumina TruSeq(R) DNA PCR-free LT kit (Cat. No. FC-121-3001). One library per lane was sequenced yielding on average 173 M read-pairs of 100bp per strain (172 – 176 M), equivalent to a 12X coverage of the mouse genome.

DNA sequence reads were aligned to the GRCm38 with alternative loci reference genome using `bwa-mem` using default parameters. Variant discovery was carried out using Platypus (Rimmer et al. 2014) where the genome of each founder sire was compared to the reference genome. Variants with quality scores less than 20, shared across two or more strains, and those present on the A/J chromosome were excluded. In addition, we analyzed our aligned files with Freebayes and GATK (Garrison and Marth 2012; DePristo et al. 2011; McKenna et al. 2010), to identify all potential variation at specific loci e.g. promoters of differentially expressed genes such as *Crem* and *Gm26448*, as these tools have a higher sensitivity to detect variants but also a higher degree of false positives (data not shown). Copy number variation analysis was carried out using CNVkit, using `wgs` method Gencode M4 annotation (Talevich et al. 2016). Our *C57BL/6J* founder male was used as a control to obtain B6 subtracted copy number ratios for all four CSS. This choice was of major importance as CNVkit does not mask X-Y pseudo-autosomal region. Thus enabling us to query copy number variation in *Mid1*. Copy number gains and losses were considered if the log2-ratio was greater than *log2*(3/2), or lower than *log2*(1/2), respectively.

Our analysis revealed 464,882 strain-specific de-novo mutations in the founder males (Rimmer et al. 2014) **(**[**Supplementary Table S2**](https://docs.google.com/spreadsheets/d/1anXtzf4XLofGKqUBhI-U4PlfijkDCzGr/edit#gid=1541101937)**,** [**Supplementary Figure S1**](https://docs.google.com/document/u/0/d/1cCM1LOEhIUw1aEJWPdr3fqvD2385_ryvmT2O96LyR3c/edit)**)**. Specifically, we identified 687, 2660, 715, and 530 non-A/J candidate strain specific de-novo single nucleotide polymorphisms (SNP) and In/Dels in the consomic strains for MMU 15, 17, 19, and X, respectively. These numbers exclude all SNP and In/Dels contained on the A/J donor chromosome of each strain, corresponding to 110,867; 24,9472; 77,824; and 21,745 for MMU 15 in B6.A-15, MMU 17 in B6.A-17, MMU 19 in B6.A-19, and MMU X in B6.A-X, respectively. Furthermore, we identified 300 de-novo mutations in the B6 founder male. Strain-specific variants were found in all chromosomes, where larger chromosomes appear to have a larger number of variants relative to smaller chromosomes. Purebred B6 had the fewest number of variants, though due to genetic drift. The Consomic strain from MMU 17 had the most variants, whereas consomic strain for B6.A-X had the least **(**[**Supplementary Table S2**](https://docs.google.com/spreadsheets/d/1anXtzf4XLofGKqUBhI-U4PlfijkDCzGr/edit#gid=1541101937)**)**.

We observed one de-novo mutation in the 3' end of the gene on the founder male for B6.A-19 (Figure 3). This gene is located on MMU X, however, given that our experimental design focused on the paternal lineage, two generations of backcrossing to purebred B6 should have replaced the MMU X that comes of the B6.A-19 founder male by one from all the B6 females in two generations. Other differentially expressed genes, such as *Gm26448* and *Crem* did not show any strain-specific variant on or upstream of its promoter within 3 Kb.

Overall copy number variation showed a simple flat profile on the majority of chromosomes across all strains, resembling B6. We identified major deterioration of MMU Y specific to the B6.A-X consomic strain ([**Supplementary Figure S1**](https://docs.google.com/document/u/0/d/1cCM1LOEhIUw1aEJWPdr3fqvD2385_ryvmT2O96LyR3c/edit)), suggesting an erosion of MMU Y on this strain (Soh et al. 2014; Case et al. 2015). However, we did not identify any differentially expressed genes on MMU Y between isogenic B6.A-X F1*^B6/B6^* to B6.C. In addition, we observed CNV regions shared across the four strains were observed in MMU 4, MMU 5, MMU 7, and MMU 12. These CNV were recurrent across several mouse strains, and have been reported by Agam et al. (2010), and Pezer et al. (2015). We considered these parts of the B6 background, and upon closer inspection identified that they fall into repetitive sequence and low complexity regions. Furthermore, the regions did not exceed our established log_2_ ratio thresholds for segment gain of log_2_(3/2) and segment loss of log_2_(1/2).

#### Figures

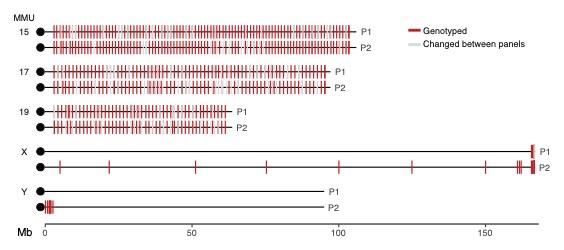

**Figure S1.** Map of SNP markers used to identify non-recombinant mice. Briefly, we selected 236 SNP markers evenly spaced throughout chromosomes 15, 17, and 19. For chromosomes X and Y we focused on those present in the XY-pseudo autosomal region. All selected markers had evidence of being polymorphic between B6 and A/J in MGI or Sanger Mouse Genomes project.

**Figure S2.** Genotypes of adult males and female/male embryo N2 backcross populations. A) Genotypes of Cohort I N2 backcross B) Genotypes of Cohort II N2 embryo backcross C) Genotypes of Cohort III N2 replicate backcross. We observe normal recombination rates in all chromosomes. Homozygous non-recombinant *B6/B6* males were found at a rate of one in every five males derived from the three autosomes. No mice observed A/J alleles in the XY-Pseudoautosomal Region.

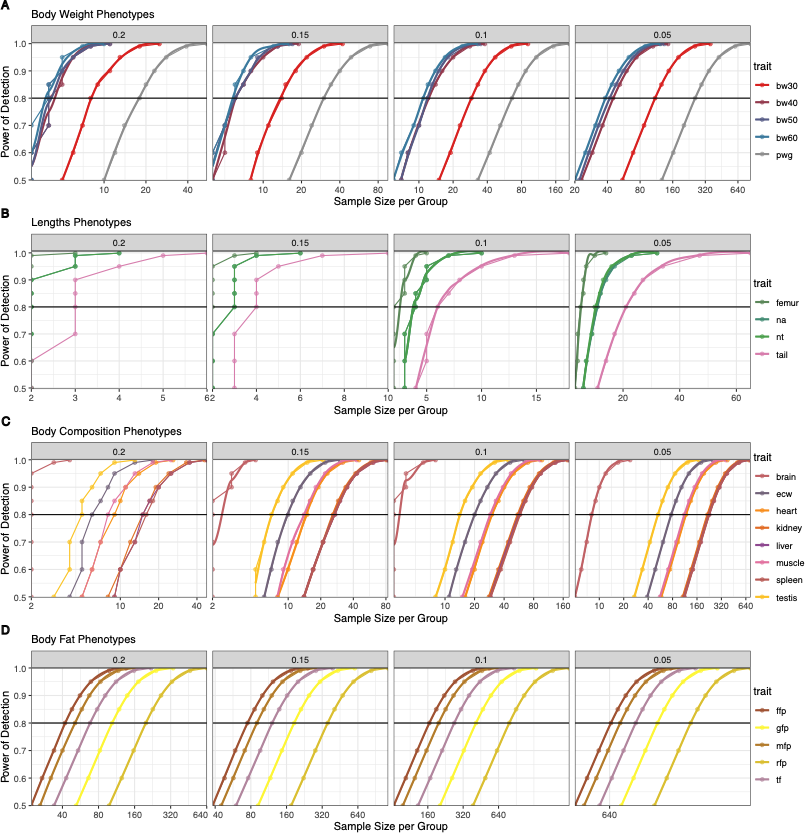
**Figure S3.** Power of detection for all anatomical phenotypes at four effect sizes, and sample size per groups. Panel columns reflect effect sizes of 20%, 15%, 10%, and 5 % of change in mean. Rows (A-D) are trait groups, **A)** Power of detection across body weight at 30, 40, 50, and 60 days, and post-weaning weight gain (30 to 60 d). **B)** Power of detection for length phenotypes. **C)** Body composition phenotypes encompassing major organs. **D)** Body fat phenotypes.

**Table S1.** RNA-seq Detection power based on Log2 Fold change across all tissues using an N = 2.

| ***Tissue*** | ***N*** | ***FC ≥ 2.0*** | ***FC ≥ 1.5*** |
| --- | --- | --- | --- |
| Pituitary | 2 | 0.97 | 0.81 |
| Liver | 2 | 0.94 | 0.37 |
| Heart | 2 | 0.96 | 0.68 |
| Embryo | 2 | 0.56 | < 0.1 |
| Placenta | 2 | 0.66 | < 0.1 |

#### References

Agam A, Yalcin B, Bhomra A, Cubin M, Webber C, Holmes C, Flint J, Mott R. 2010. Elusive copy number variation in the mouse genome. *PLoS ONE* **5**: e12839.

Case LK, Wall EH, Osmanski EE, Dragon JA, Saligrama N, Zachary JF, Lemos B, Blankenhorn EP, Teuscher C. 2015. Copy number variation in Y chromosome multicopy genes is linked to a paternal parent-of-origin effect on CNS autoimmune disease in female offspring. *Genome Biology* **16**: 28-14.

Cox A, Ackert-Bicknell CL, Dumont BL, Ding Y, Bell JT, Brockmann GA, Wergedal JE, Bult C, Paigen B, Flint J et al. 2009. A New Standard Genetic Map for the Laboratory Mouse. *Genetics* **182**: 1335-1344.

Nadeau JH, Singer JB, Matin A, Lander ES. 2000. Analysing complex genetic traits with chromosome substitution strains. *Nature Genetics* **24**: 221-225.

Pezer Ž, Harr B, Teschke M, Babiker H, Tautz D. 2015. Divergence patterns of genic copy number variation in natural populations of the house mouse (Mus musculus domesticus) reveal three conserved genes with major population-specific expansions. *Genome Research* **25**: 1114-1124.

Rimmer A, Phan H, Mathieson I, Iqbal Z, Twigg SRF, Consortium W, Wilkie AOM, McVean G, Lunter G. 2014. Integrating mapping-, assembly- and haplotype-based approaches for calling variants in clinical sequencing applications. *Nature Genetics* **46**: 912-918.

Soh YQS, Alföldi J, Pyntikova T, Brown LG, Graves T, Minx PJ, Fulton RS, Kremitzki C, Koutseva N, Mueller JL et al. 2014. Sequencing the Mouse Y Chromosome Reveals Convergent Gene Acquisition and Amplification on Both Sex Chromosomes. *Cell* **159**: 800-813.

Talevich E, Shain AH, Botton T, Bastian BC. 2016. CNVkit: Genome-Wide Copy Number Detection and Visualization from Targeted DNA Sequencing. *PLoS Computational Biology* **12**: e1004873.
