## Supplemental Figures for "Lack of evidence supporting transgenerational effects of non-transmitted paternal alleles on the murine transcriptome"

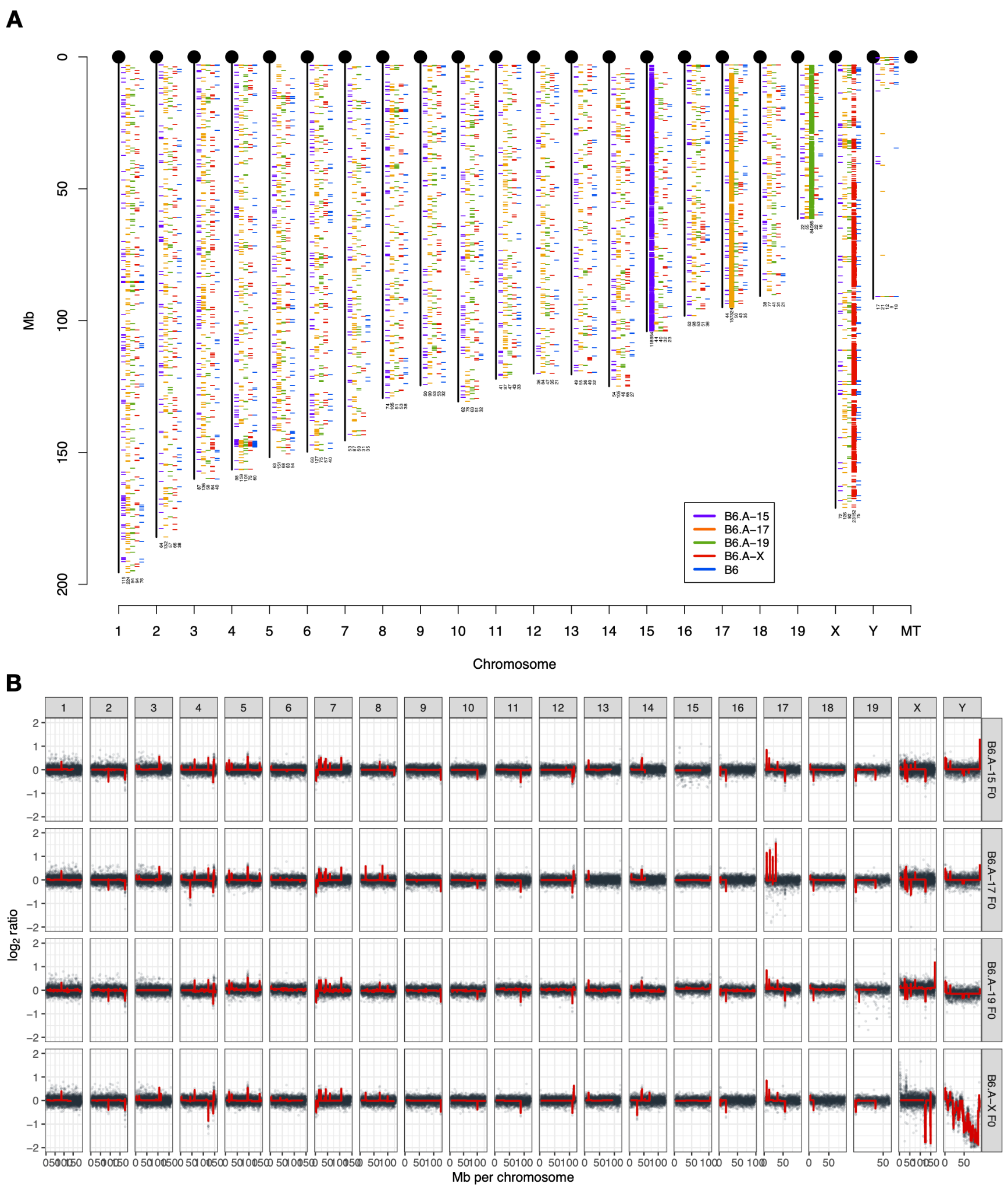
**Supplementary Figure S1.** The mutational and copy number landscapes of de-novo and A/J alleles across the B6.A-<N> founder males of the tested CSS. A) Chromosome number is along the x-axis and Mb coordinate along the y-axis. Each tick mark represents a heterozygous de-novo mutation, and the corresponding number of mutations is listed at the end of each chromosome. The figure shows 16 loci of shared heterozygosity across these strains, however, the individual mutations in these loci are private to each CSS. As expected, MMU 15, 17, 19, and X show enrichment of mutations for their corresponding CSS. B) Copy number variation across the B6.A-<N> founder males of the tested CSS. Black points show the *log2* ratio of individual bins, whereas the red line reflects the segment mean log2 ratio. Each track represents the copy number profile of each CSS founder male normalized to the C57BL/6J founder.

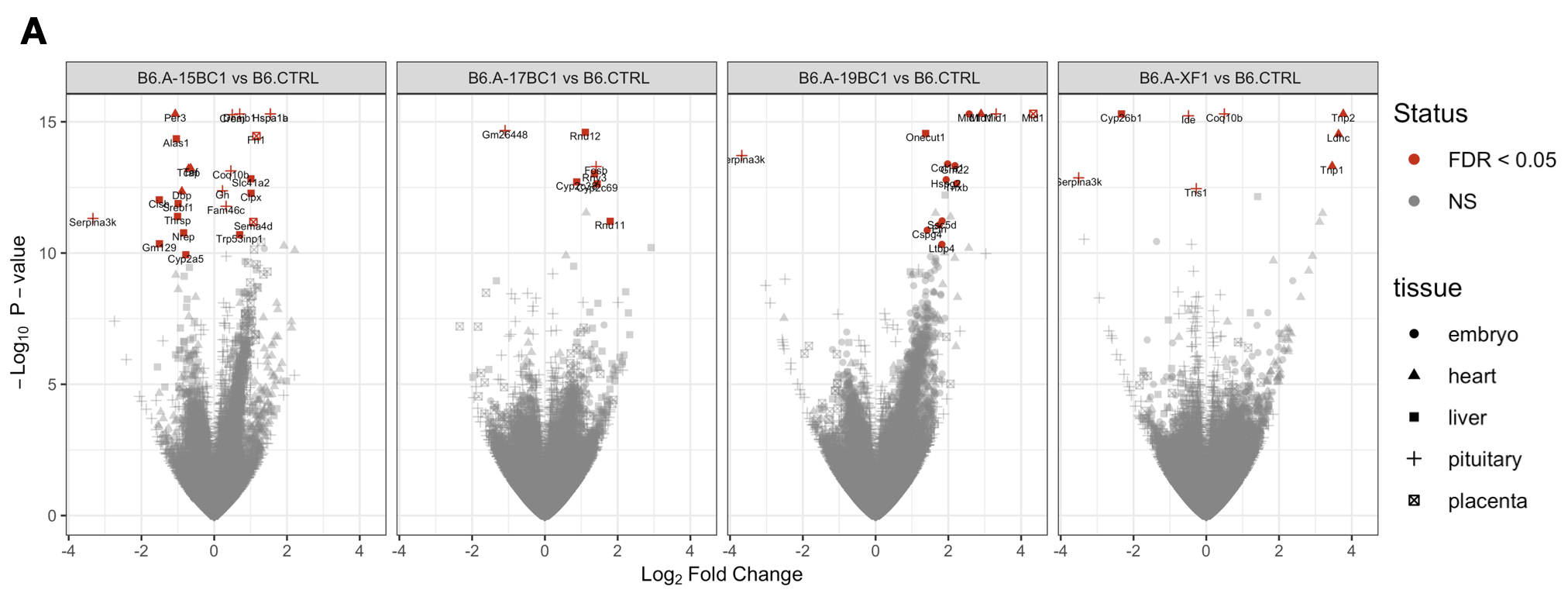

**Supplementary Figure S2.** Volcano plots of differential expression across all derived isogenic N2 mice from the four CSS. Significant differentially expressed genes (FDR ≤ 0.05) are marked in red, labeled, and point shape coded for each tissue.

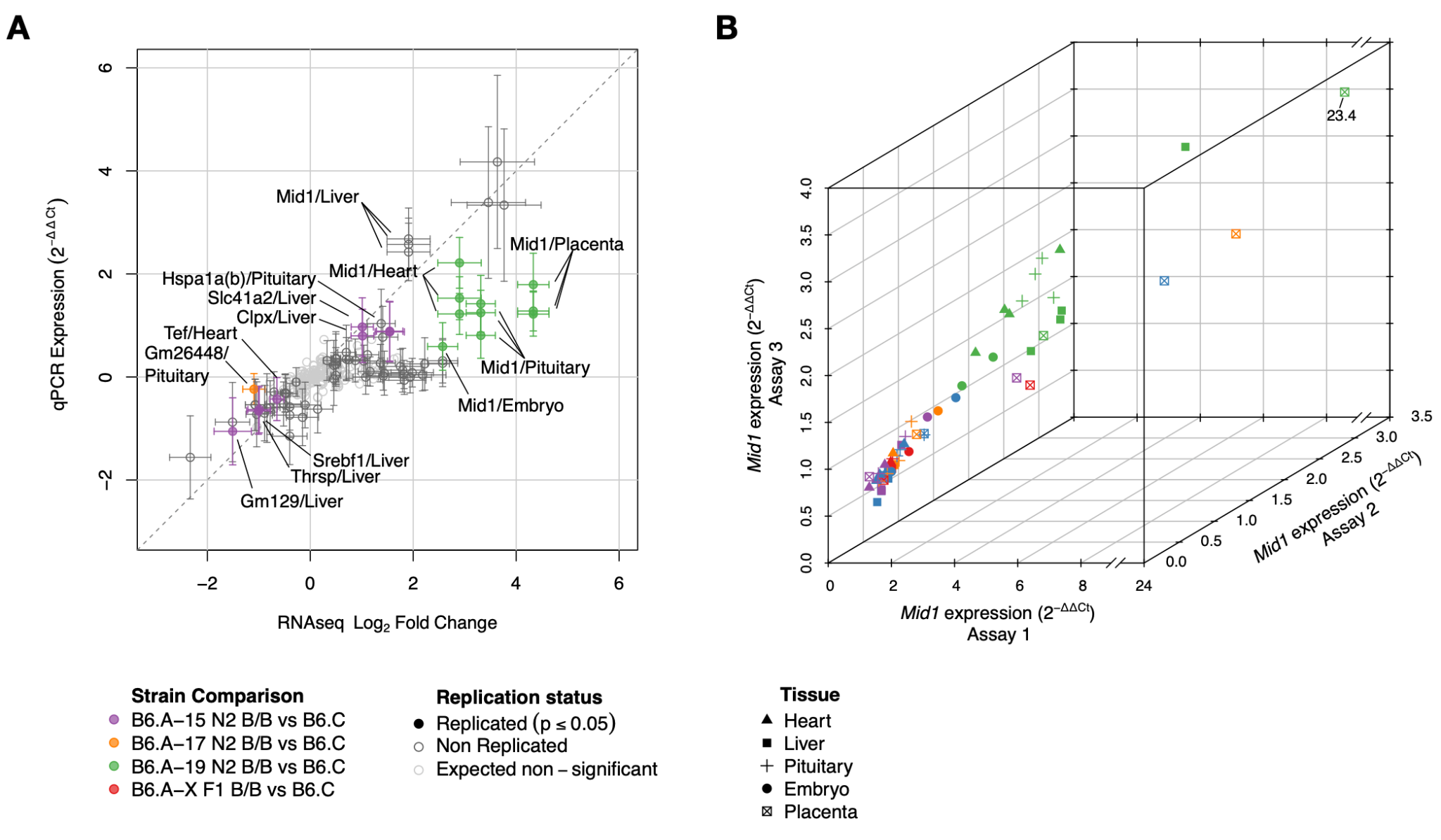

**Supplementary Figure S3.** Global expression and *Mid1* effect comparisons across replicates. **A)** Cross-platform replication between RNA-seq Log_2_ Fold Change and qPCR Expression (as 2^-ΔΔCt^) showing persistent differential expression in 9 genes, including *Mid1* from the B6.A-19 lineage. **B)** Multidimensional correlation of *Mid1* gene expression across 3 assays measuring distinct *Mid1* isoforms show a clear trend of over-expression in B6-A-19 N2 *^B6/B6^* when compared to B6 contemporary group mice across all tissues. This effect is not evident in isogenic mice from the other strains, except in the placenta where it has a functional role.

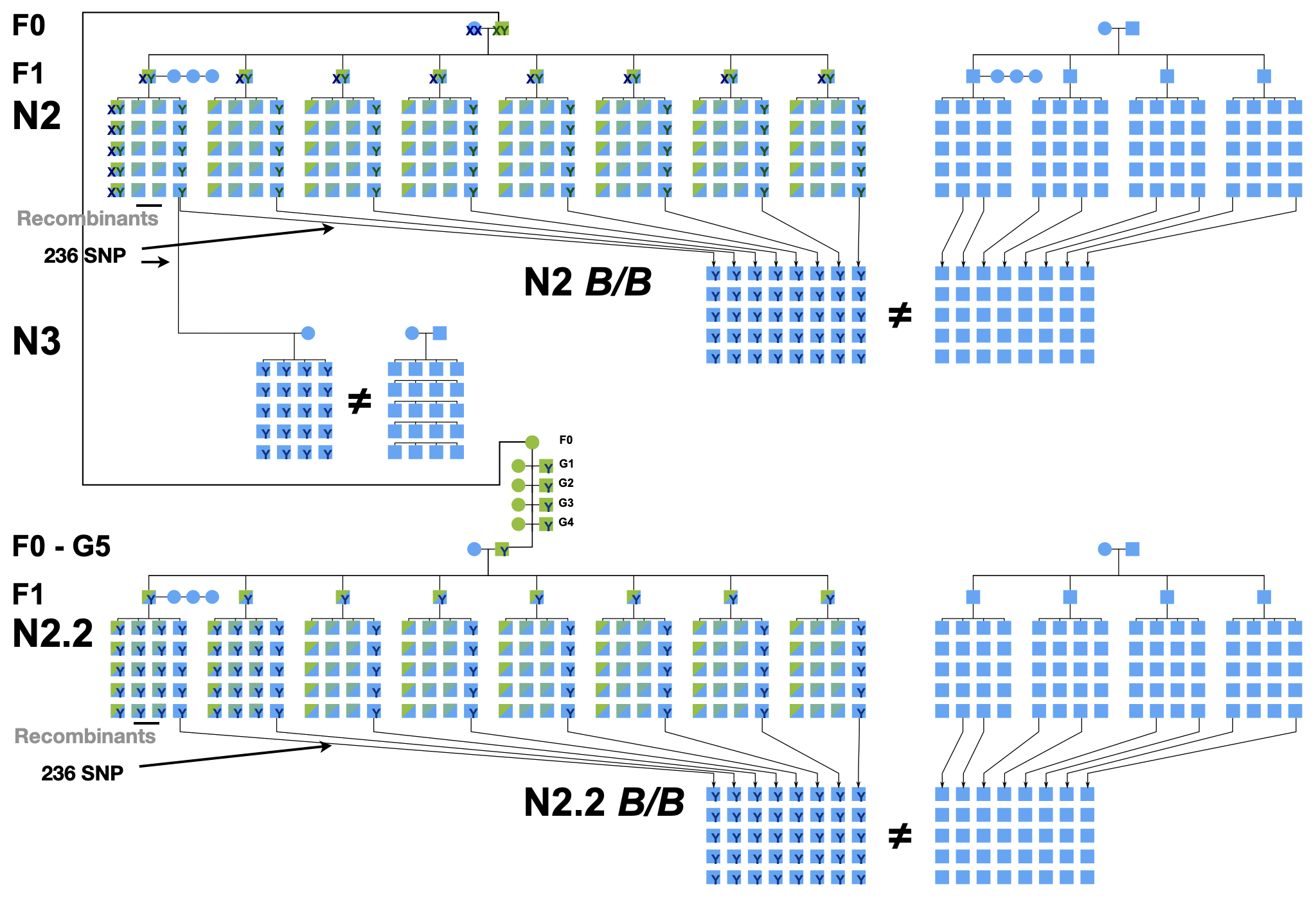

**Supplementary Figure S4.** Inheritance model of the *Mid1* segmental duplication across the B6-A-19 male lineage throughout the discovery, validation, and persistence experimental crosses. Pedigree initiates with a single founder B6.A-19 F0 sire mated to a single C57BL/6J female. Male offspring in the F1 inherit the MMU Y from their sire and MMU X from the dam. Subsequent backcrosses were all from male F1 heterozygous to C57BL/6J females, preserving MMU Y lineage from the founder male. In our case, all N2 backcrossed males would inherit the grand-sire MMU Y, and grand-dam X. In the last backcross N3, males would continue to inherit the sire lineage of MMU Y. The B6.A-19 CSS was preserved for five generations through brother-sister mating. The replicate backcross was initiated with a G5 B6.A-19 male mated to a C57BL/6J female, and the resulting F1 backcrossed to C57BL/6J females to generate the N2. Given the structure of the cross, the MMU Y was preserved throughout the male lineage and MMU X through the female lineage.

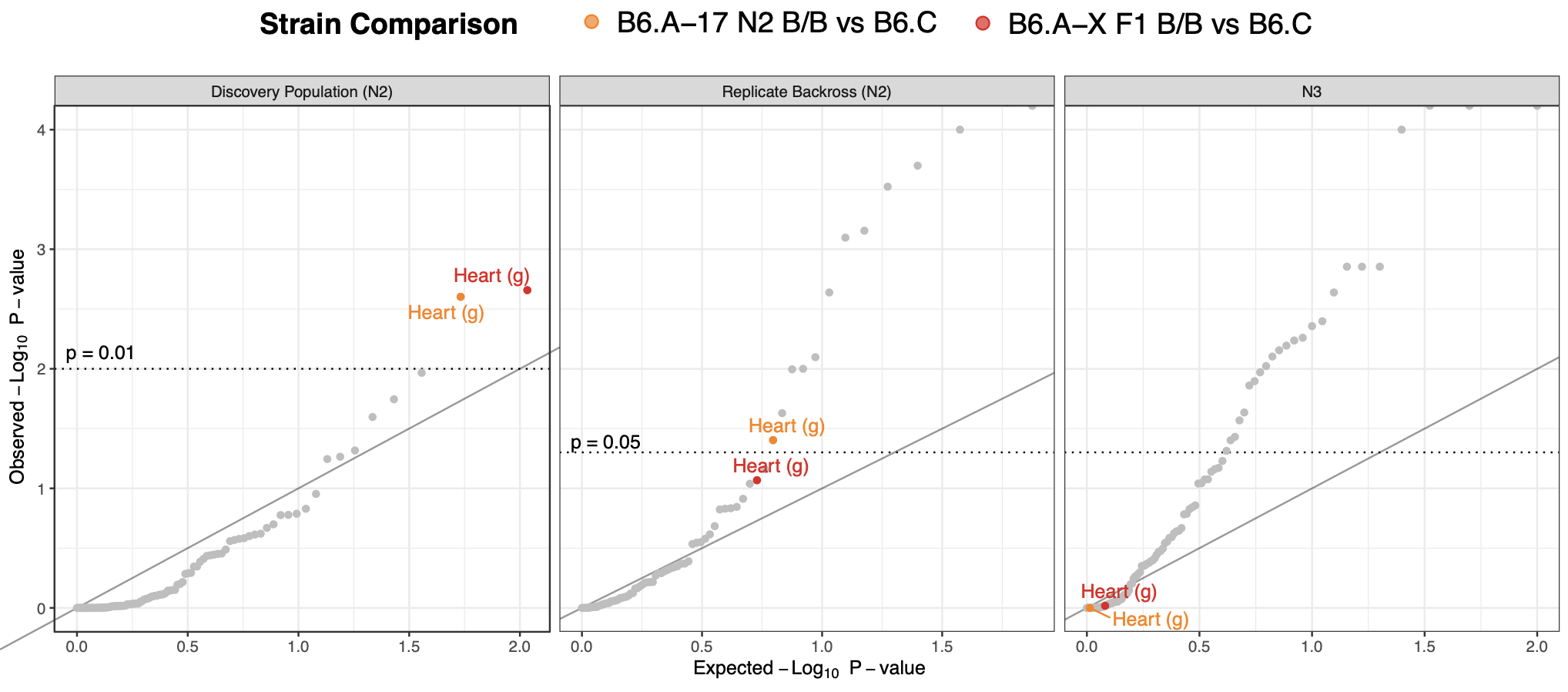

**Supplementary Figure S5.** QQ-plots of -log_10_ *p*-values of all 25 anatomical traits across all three analyzed cohorts. (Left panel) Initial results suggest one traits in two strains Heart weight to be significantly different between in B6.A-17 N2 *^B6/B6^* and B6.A-X F1 *^B6/B6^* isogenic relative to purebred B6 control mice (p ≤ 0.01). (middle panel) Replication of the experiment with an independent N2 cross shows only Heart weight in B6.A-17 N2 *^B6/B6^* continued to be higher than B6 control mice (p < 0.05, one-tail test). (right panel) This effect is not reproducible in the next generation of successive backcrossing (N3). All other traits shown in gray were not significantly different in the first generation and were not further analyzed in subsequent generations.

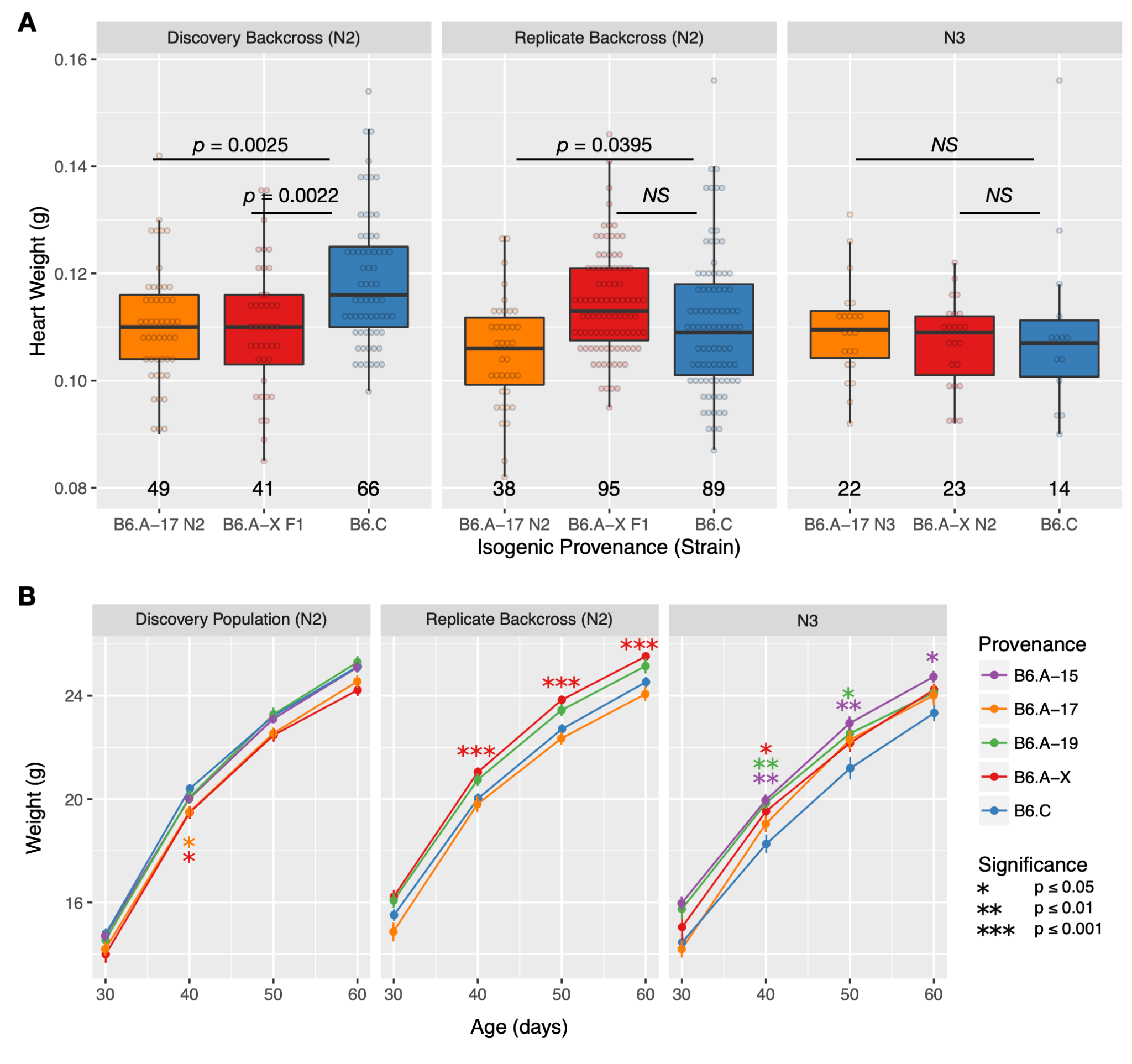

**Supplementary Figure S6.** Heart, and body weights weight across the Discovery (N2), Replicate (N2), and Persistence (N3) backcrosses in B6.A-17 N2 *^B6/B6^*, B6.A-X F1 *^B6/B6^*, and B6 Control mice. **A)** A lower heart weight is observed between B6.A-17 N2 *^B6/B6^* and B6.C mice in both Discovery and Replicate N2 backcrosses, but not in the Persistence (N3) backcross. The effect on B6.A-X F1 *^B6/B6^* and B6 Control mice was only observed in the Discovery N2 cross. These results suggest transgenerational genetic effects are not carried over beyond a single-backcross. **B)** Growth curves of isogenic B6.A-<N> N2 *^B6/B6^* male mice. Each panel represents a distinct population, with age in days on the x-axis and body weight in grams on the y-axis. Each dot represents the mean body weight of the population ± 1 SD, color-coded for the founder strain (provenance) of the isogenic B6.A-N N2 *^B6/B6^* mice. Significance asterisks represent the comparison of a B6.A-<N> N2 *^B6/B6^* male mice to their B6.C contemporary group and are also color-coded to show which isogenic comparisons were significant. In the discovery population, mice from the B6.A-17 and B6.A-X lineage were significantly lower in body weight at 40 days when compared to B6.C. In the replicate backcross, we observed an opposite effect, where B6.A-X has higher body weights than the B6.C, and B6.A-17 does not show any significant differences at any time-point. The N3 shows additional changes where other isogenic lineages show distinct body weight changes albeit, specifically at 40, 50, and 60 days of age for the B6.A-15 lineage. Data is available in Supplementary Table S2.
